# The heme oxygenase-1 metalloporphyrin inhibitor stannsoporfin enhances the bactericidal activity of a novel regimen for multidrug-resistant tuberculosis in a murine model

**DOI:** 10.1101/2023.08.09.552716

**Authors:** Jennie Ruelas Castillo, Pranita Neupane, Styliani Karanika, Stefanie Krug, Darla Quijada, Andrew Garcia, Samuel Ayeh, Addis Yilma, Diego L. Costa, Alan Sher, Nader Fotouhi, Natalya Serbina, Petros C. Karakousis

## Abstract

Multidrug-resistant (MDR) *Mycobacterium tuberculosis* (Mtb) poses significant challenges to global tuberculosis (TB) control efforts. Host-directed therapies (HDT) offer a novel approach for TB treatment by enhancing immune-mediated clearance of Mtb. Prior preclinical studies found that inhibition of heme oxygenase-1 (HO-1), an enzyme involved in heme metabolism, with tin-protoporphyrin IX (SnPP) significantly reduced mouse lung bacillary burden when co-administered with the first-line antitubercular regimen. Here we evaluated the adjunctive HDT activity of a novel HO-1 inhibitor, stannsoporfin (SnMP), in combination with a novel MDR-TB regimen comprising a next-generation diarylquinoline, TBAJ-876 (S), pretomanid (Pa), and a new oxazolidinone, TBI-223 (O) (collectively, SPaO) in Mtb-infected BALB/c mice. After 4 weeks of treatment, SPaO + SnMP 5 mg/kg reduced mean lung bacillary burden by an additional 0.69 log_10_ (P=0.01) relative to SPaO alone. As early as 2 weeks post-treatment initiation, SnMP adjunctive therapy differentially altered the expression of pro-inflammatory cytokine genes, and CD38, a marker of M1 macrophages. Next, we evaluated the sterilizing potential of SnMP adjunctive therapy in a mouse model of microbiological relapse. After 6 weeks of treatment, SPaO + SnMP 10 mg/kg reduced lung bacterial burdens to 0.71 ± 0.23 log_10_ CFU, a 0.78 log-fold greater decrease in lung CFU compared to SpaO alone (P=0.005). However, adjunctive SnMP did not reduce microbiological relapse rates after 5 or 6 weeks of treatment. SnMP was well tolerated and did not significantly alter gross or histological lung pathology. SnMP is a promising HDT candidate requiring further study in combination with regimens for drug-resistant TB.

## Introduction

*Mycobacterium tuberculosis* (Mtb) is the causative agent of tuberculosis (TB), a deadly disease that claims millions of lives yearly. The emergence of multidrug-resistant TB (MDR TB), which is resistant to at least two of the most potent first-line drugs, rifampin, and isoniazid, is a significant challenge to global health [1]], [2]]. The World Health Organization estimated that in 2021, there were 465,000 new cases of MDR TB worldwide, and only 60% of these cases were treated successfully [1]. MDR-TB treatment is lengthy, complex, expensive, and requires the use of second-line drugs that are less effective and more toxic, highlighting the urgent need for novel therapeutics [3], [4]. Given the global burden of MDR TB, host-directed therapies (HDT) targeted at boosting the immune system have the potential to contribute to tuberculosis control to prevent further cross-resistance with current antibiotics [5], [6].

Heme oxygenase-1 (HO-1) is an enzyme involved in the degradation of heme into biliverdin, carbon monoxide (CO), and iron [7]–[10]. HO-1 is a known biomarker for active TB in humans [11]. HO-1 also plays a crucial role in the survival of Mtb inside the host by suppressing immune responses and promoting bacterial growth [12]. Several studies have shown that Mtb infection induces HO-1 expression in macrophages [8], [13], leading to the generation of CO, which has immunomodulatory properties and can suppress the production of pro-inflammatory cytokines, such as TNF-α, IL-1β, and IL-6 [14], [15]. Additionally, the production of CO can inhibit the activity of T cells and natural killer cells, which are essential components of the host immune response against Mtb [16]. HO-1 induction can drive macrophage polarization towards an anti-inflammatory, M2 phenotype, which is associated with reduced microbicidal activity [17], [18] and increased persistence of Mtb in the host [19].

Several studies have highlighted the potential utility of HO-1 inhibitors as HDT for TB. Costa *et al.* showed previously that pharmacological inhibition of HO-1 with the metalloporphyrin tin protoporphyrin (SnPP) during the chronic phase of infection in the C57BL/6 murine model of TB reduced the lung bacillary burden alone and also augmented the activity of the standard regimen against drug-susceptible TB [20]. HO-1 inhibition promoted the differentiation of CD4^+^ T cells into interferon gamma (IFNγ)-producing Th1 cells. Additionally, SnPP enhanced IFN-γ-dependent, nitric oxide synthase 2 (NOS2)-induced nitric oxide production by macrophages, resulting in enhanced control of bacterial growth [21]. Another compound in this class, the HO-1 inhibitor stannsoporfin (SnMP), has been in clinical development as a therapy for hyperbilirubinemia in neonates [22]. SnMP has been shown to induce the activation, proliferation, and maturation of naïve CD4^+^ and CD8^+^ T cells via interactions with CD14^+^ monocytes in *vitro* [23].

In the current study, we compared the adjunctive bactericidal activity of SnPP and SnMP when co-administered with a novel MDR-TB regimen containing TBAJ-876 (S), pretomanid (Pa), and TBI-223 (O); (collectively, SPaO) or the first-line drug regimen rifampin (R), isoniazid (H), and pyrazinamide (Z); (collectively, RHZ) against chronic lung infection with drug-susceptible Mtb in BALB/c mice. In addition, we tested the ability of adjunctive SnMP to shorten the duration of curative treatment in a murine model of microbiological relapse.

## Results

### In chronically infected BALB/c mice, adjunctive therapy with SnMP increases the bactericidal activity of SPaO

We investigated SnPP and SnMP as adjunctive HDT agents in BALB/c mice infected with Mtb H37Rv (Fig 1A). Each compound was given alone or co-administered with human-equivalent doses of the first-line regimen RHZ for a total of 6 weeks or the enhanced-potency MDR regimen SPaO for a total of 4 weeks. In a previous study [20], SnPP was dosed at 5 mg/kg, but, in the current study, the dose was increased to 10 mg/kg to more closely approximate the drug exposures observed with SnMP 5 mg/kg. Briefly, plasma pharmacokinetics profiles were determined for SnPP and SnMP up to 24 hours following drug administration. The last time point with detectable blood concentrations was after 8 hours. SnPP 10 mg/kg and SnMP 5 mg/kg achieved similar 24-hour drug exposures in plasma (Fig S1), supporting the selection of these doses for the primary study. Over the course of the study, vehicle-treated animals maintained a steady lung colony-forming units (CFU) plateau, while each of the other regimens reduced the mean lung CFU burden steadily over time (Fig 1B). After 2 weeks of treatment (Fig 1C, Table1), monotherapy with SnPP 10 mg/kg or SnMP 5 mg/kg reduced the mean lung bacillary burden by 0.88 log_10_ (P = 0.0001) and 0.67 log_10_ CFU (P = 0.0004), respectively, relative to the vehicle control. Similarly, after 4 and 6 weeks of treatment, SnPP 10 mg/kg reduced the mean lung bacterial burden by 1.2 log_10_ CFU (P = 0.0001) and 1.1 log_10_ CFU (P = 0.0001), respectively, relative to the vehicle control (Figs 1D, 1E). Although SnMP 5 mg/kg significantly reduced the mean lung bacterial burden by 1.1 log_10_ CFU (P = 0.001; Fig 1D) relative to the vehicle control after 4 weeks of treatment, this bactericidal effect was not sustained at 6 weeks (0.4 log_10_ CFU, P = 0.09; Fig 1E). After 2 weeks of treatment, neither SnPP nor SnMP adjunctive therapy showed an additive effect when co-administered with SPaO (Fig 1C). However, after 4 weeks of treatment (Fig 1D), SPaO + SnMP 5 mg/kg reduced the mean lung bacillary burden to 2.49 ± 0.12 log_10_ CFU, representing an additional 0.69 log_10_ CFU reduction compared to SPaO alone (P = 0.01). Adjunctive treatment with SnMP 5 mg/kg or SnPP 10 mg/kg did not significantly enhance the bactericidal activity of RHZ at 4 weeks (P = 0.52 and P = 0.89, respectively; Fig 1D) or 6 weeks (P = 0.97 and P = 0.86, respectively; Fig 1E).

**Figure 1:**
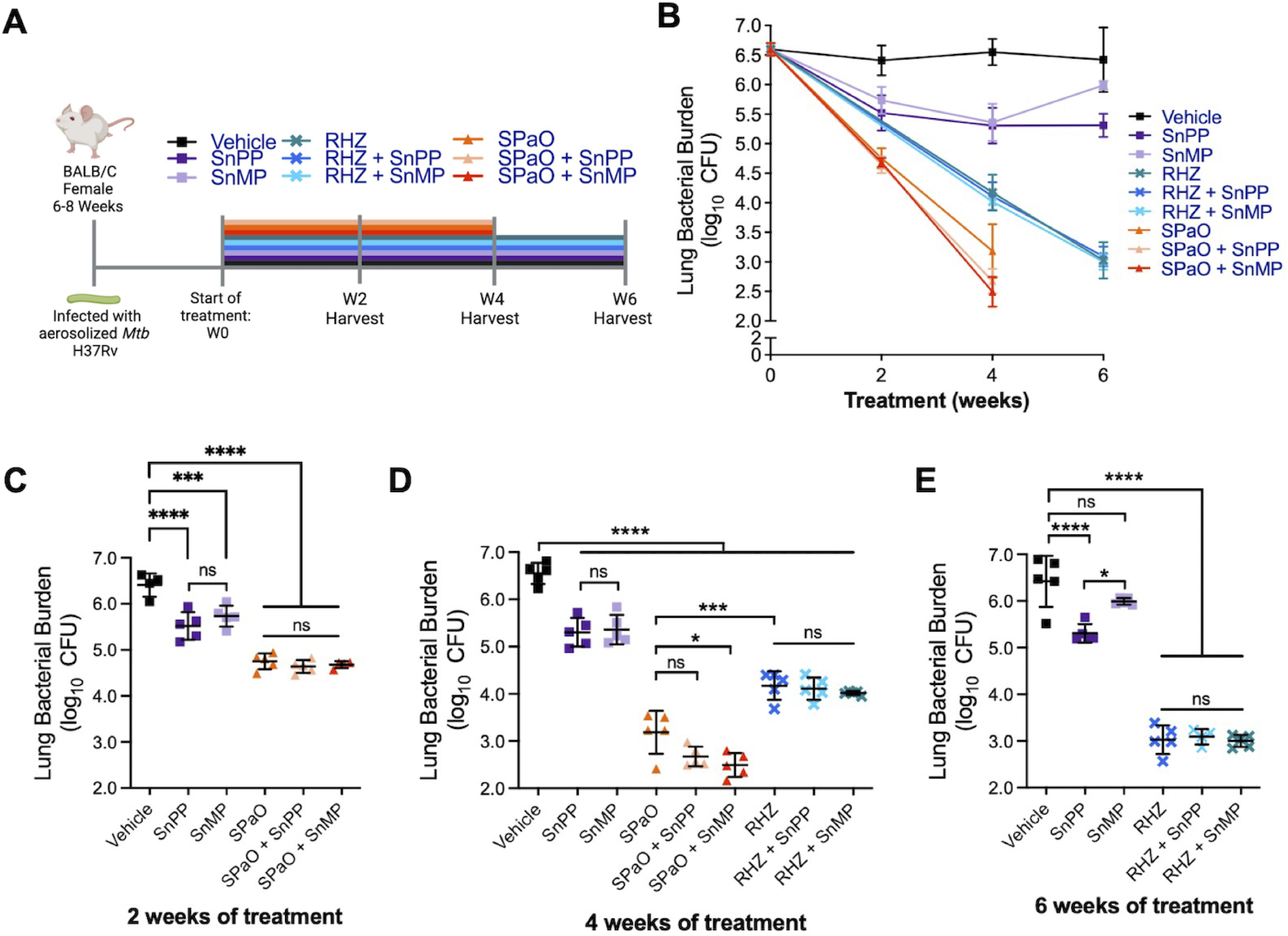
HO-1 inhibitors increase the antitubercular activity of a novel MDR regimen in a mouse model of chronic TB. A) Experimental design of mouse experiments. Each line represents a different treatment group and length of treatment. BALB/c mice were aerosol-infected with ∼100 bacilli of Mtb H37Rv. Treatment was initiated 4 weeks after aerosol infection. B) CFU means ± standard deviation (SD) of lung bacillary burden after treatment initiation (week 0). Scatterplot of lung mycobacterial burden after 2 weeks C), 4 weeks D) or 6 weeks E) of treatment. Each point represents the CFU per mouse lung from individual, the error bars represent the mean ± SD. RHZ: rifampin/isoniazid/pyrazinamide; SPaO: TBAJ-876/pretomanid/TBI-223; SnPP: tin-protoporphyrin IX; SnMP: stannsoporfin; CFU: colony-forming units. For panels B-E, each experimental group consisted of four to five mice. For panels C-E, statistical analysis was performed by an ordinary one-way ANOVA followed by Dunnett’s correction to control for multiple comparisons and Bonferroni correction for treatment groups with the same antibiotic backbone. * = P < 0.05; ** = P < 0.01; *** = P < 0.001; **** = P < 0.0001; ns: not significant.

### HO-1 inhibition does not exacerbate lung inflammation and regulates pro-inflammatory immune responses

The chronic stages of TB disease are characterized by severe lung inflammation, which results in long-term lung dysfunction in one quarter of cases despite microbiological cure [24], [25]. After 4-6 weeks of infection, the gross pulmonary pathology of Mtb-infected BALB/c mice is characterized by pronounced tubercular lesions. At the histopathological level, macrophage and lymphocyte inflammatory aggregates fill the alveolar spaces, representing cellular granulomatous lesions [26]. HO-1^–/–^ knock-out mice develop a progressive inflammatory state [13], [27], and splenocytes isolated from these mice respond to lipopolysaccharide stimulation by producing inflammatory cytokines [4], [13]. Chronic treatment with SnPP in wild-type mice can induce tubulointerstitial inflammation and fibrosis [28], while SnMP induces the expression of pro-inflammatory cytokines in human peripheral blood mononuclear cells [29]. Robust expression of pro-inflammatory cytokines can exacerbate lung inflammation, however, when controlled, it can offer antimycobacterial protection to the host [30].

To determine whether HO-1 inhibition modulates TB-associated lung pathology, we first analyzed the lung weight to body weight ratio (lung/body weight) of Mtb-infected mice treated with SnPP or SnMP alone or as adjunctive HDT with RHZ or SPaO as a gross indicator of lung inflammation. An increase in this ratio reflects an increase in lung inflammation caused by cellular infiltration and the presence of tissue lesions consistent with disease exacerbation [26], [31]. Antitubercular treatment with RHZ for 4 or 6 weeks or with SPaO for 2 or 4 weeks significantly reduced (P < 0.0001) the mean lung/body weight ratio relative to vehicle (Fig 2A-C). SnMP or SnPP monotherapy for 2 or 6 weeks also significantly reduced the lung/body weight ratio relative to vehicle (Fig 2A, C), although the decrease in this ratio was not statistically significant after 4 weeks of treatment (Fig 2B). The lung/body weight ratio was significantly (P < 0.0001) correlated with the lung CFU burden in each of the treatment groups, as assessed by the Pearson correlation coefficient (r = 0.9074 after 2 weeks of treatment; r = 0.8251 after 4 weeks of treatment; and r = 0.9282 after 6 weeks of treatment) (Fig S2). Grossly, after 4 weeks of treatment, the lungs of mice treated with vehicle, SnPP or SnMP contained more pronounced tubercle lesions compared to those of mice receiving the antitubercular drug regimens (Fig 2D).

**Figure 2:**
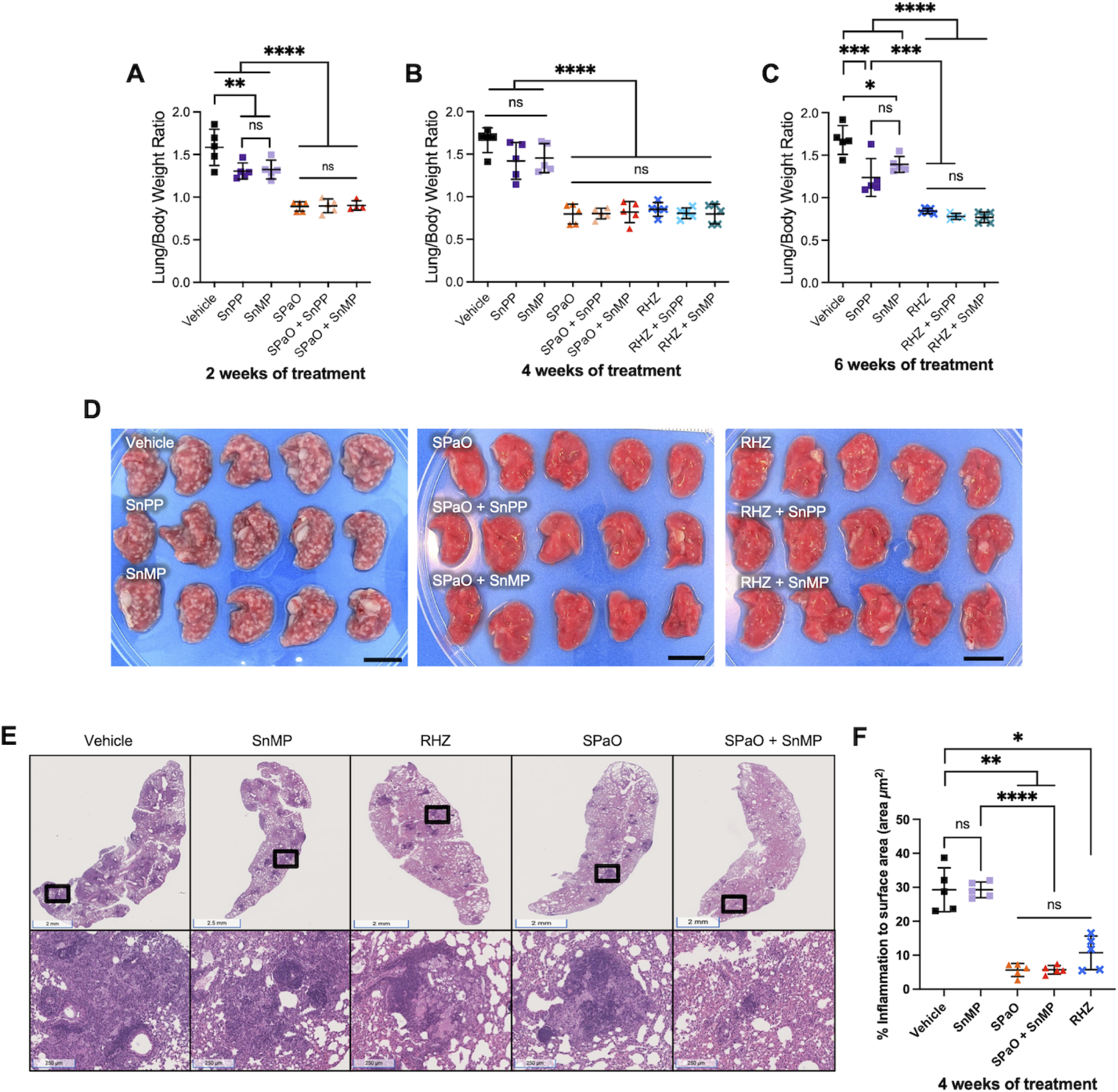
SnMP adjunctive therapy does not exacerbate gross lung inflammation. A, B, C) Scatterplot of lung/body weight ratio. D) Image of the right lung at time of harvest after 4 weeks of treatment. Scale bar = 1 cm. E) Hematoxylin and eosin staining of left lung medial apex to base sections demonstrating the histopathology of different treatment groups after 4 weeks of treatment. Top row are representative histopathology images of the groups. Bottom row are areas of inflammation within the images above (magnification of the black boxes above). F) Quantitative analysis of histopathology after 4 weeks of treatment for selected groups. g: grams; RHZ: rifampin/isoniazid/pyrazinamide; SPaO: TBAJ-876 (S)/pretomanid (Pa)/TBI-223 (O); SnPP: tin-protoporphyrin IX; SnMP: stannsoporfin. For panels A-C and F, each point represents data from individual mice, each experimental group consisted of four to five mice, and the bar represents the means ± SD. Statistical analysis was performed by an ordinary one-way ANOVA followed by Tukeys multiple comparison test for panels A-C and for panel F, Welch’s ANOVA test was conducted. ns: not significant; ** = P < 0.001; *** = P < 0.001; **** = P < 0.0001.

Given the adjunctive activity of SnMP with SPaO after 4 weeks of treatment, we next quantified the percentage of lung surface area (μm^2^) affected by inflammation at this time point using hematoxylin and eosin staining to determine if adjunctive SnMP had any impact on macrophage and lymphocytic infiltration in TB lung lesions (Fig 2E). The predominant inflammatory lesions observed were cellular granulomas containing myeloid cores and lymphocytic cuffs. As expected, the percentage of lung surface area involved by inflammation was significantly reduced by the antitubercular regimens RHZ (10.71% ± 4.9), SPaO (5.65% ± 1.9) relative to the vehicle control (29.26% ± 6.4; P = 0.011, and P = 0.003 respectively). However, there was no significant difference in percentage of lung surface area involved by inflammation between vehicle (29.26% ± 6.4) and SnMP monotherapy (29.26% ± 2.3) (P > 0.99). Similarly, SnMP adjunctive therapy did not significantly reduce the percentage of lung surface area inflammation relative to RHZ alone (10.71% ± 4.9 vs 5.74 + 1.2, P = 0.41) or SPaO alone (5.65% ± 1.9 vs 5.74 + 1.2, P > 0.99).

In order to further investigate the adjunctive contribution of SnMP to the bactericidal activity of SPaO observed at 4 weeks of treatment (Table 1), we used RT-qPCR to evaluate the expression of genes encoding pro-inflammatory cytokines and macrophage markers in lung homogenates at week 2, at which time a significant difference in lung bacillary burden was observed between treatment groups. Interferon-gamma (IFN-γ) and tumor necrosis factor-alpha (TNF-α) are important macrophage-activating cytokines [32], [33]. NOS2 expression in macrophages is upregulated in response to Mtb infection and IFN-γ activation is required for successful bacterial load reduction following pharmacological inhibition of HO-1 [21]. SnMP monotherapy was associated with reduced expression of *Tnf-α* (P = 0.005, Fig 3C) relative to vehicle. SPaO alone was associated with a further reduction in expression of *Ifn-γ* (P = 0.002, Fig 3B), and *Tnf-α* (P = 0.001, Fig 3C); while SPaO + SnMP showed reduced expression of *Nos2* (P = 0.03, Fig 3A), *Ifn-γ* (P = 0.006, Fig 3B), and *Tnf-α* (P = 0.03, Fig 3C) relative to vehicle. These results are consistent with decreased cytokine production in response to reduced mean lung bacterial burdens in these groups. *Nos2, Ifn-γ,* and *Tnf-α* expression was comparable in the lungs of mice treated with SPaO or SPaO + SnMP (Fig 3A-C). Next, we looked for expression changes in the mouse macrophage markers CD38 and early growth response protein 2 (EGR2), a transcriptional regulator/M1 marker and a negative T-cell regulator/M2 marker, respectively [34], [35]. SnMP monotherapy, but not SnMP adjunctive therapy in combination with SPaO, was associated with significantly higher expression of *Cd38* (P < 0.0001) compared to any other treatment group (Fig 3D). In contrast, *Egr2* expression (Fig 3E) did not differ significantly among groups. Overall, these data are consistent with reduced expression of pro-inflammatory cytokine genes as a result of reduced mean lung CFU in the different treatment groups as compared to the vehicle group (Table 1). However, SnMP treatment also appeared to induce the expression of *Cd38*, potentially polarizing lung macrophages to an M1 phenotype, which is more effective in killing intracellular Mtb.

**Figure 3:**
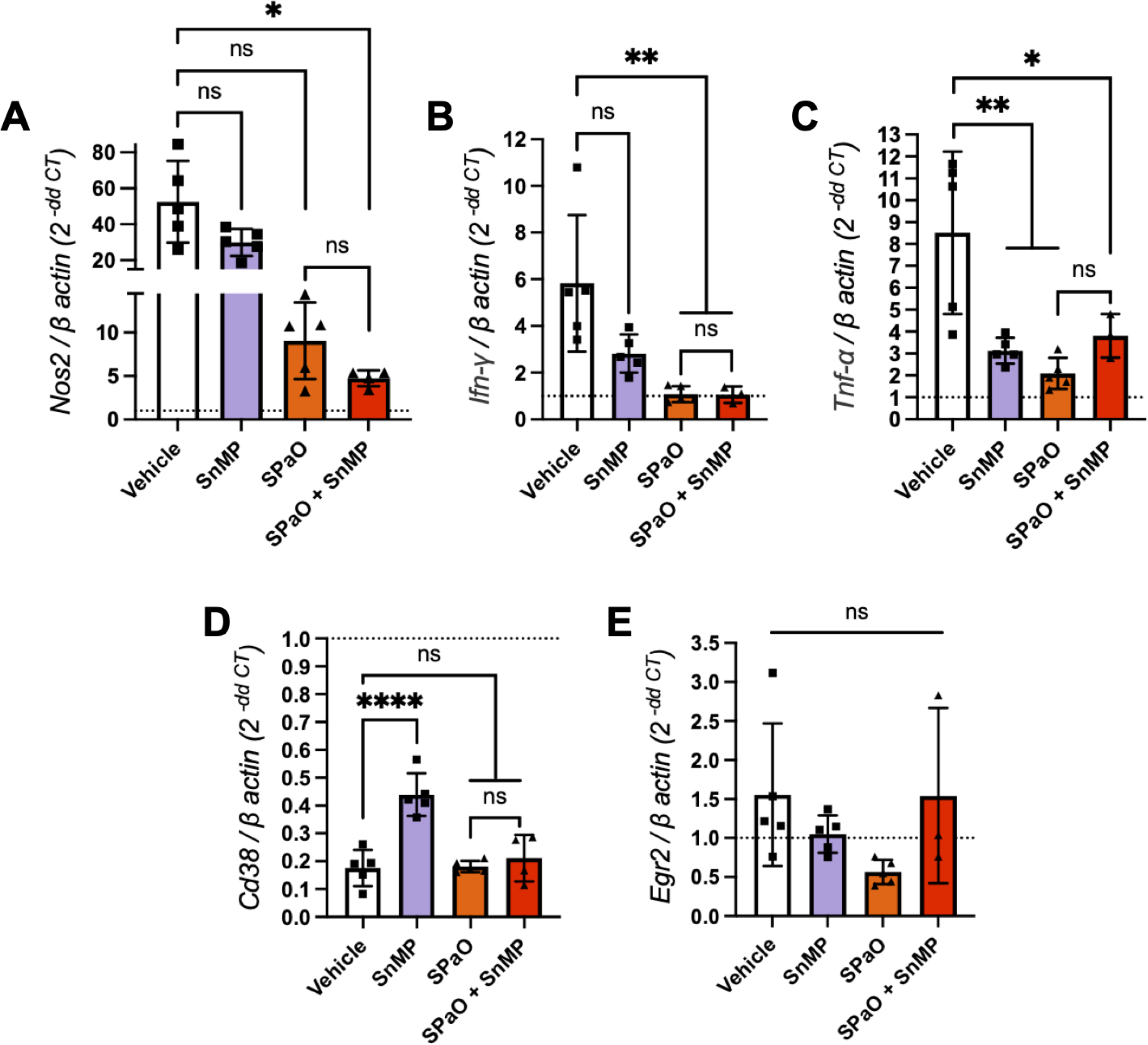
Genes encoding M1 macrophage markers were upregulated in the lungs after two weeks of treatment with SnMP. RT-cPCR was performed for the expression of the following genes in snap-frozen lung homogenates: A) *Nos2*; B) *Ifn-γ*; C) *Tnf-α*; D) *Cd38*; E) *Egr2*. The dotted line represents the uninfected normalization of mRNA expression, SPaO: TBAJ-876 (S)/Pretomanid (Pa)/TBI-223 (O); SnMP: stannsoporfin. Each bar represents a treatment group consisting of three to five mice. Data represent individual data points with means ± SD of the results. Panel A was analysed by the Welch’s ANOVA test while panels B-E were tested by an ordinary one-way ANOVA followed by Tukeys multiple comparison test. ns: not significant. * = P < 0.05; ** = P < 0.001; *** = P < 0.001, **** = P < 0.0001.

**Table 1:** Lung CFU count for time course and relapse study. dpi = day post infection; W0 = week 0 (start of treatment); W2 = Week 2; W4 = Week 4; W5 = Week 5; W6 = Week 6

| Regimen | Mean lung log<sub>10</sub> CFU count (±SD) at: |  |  |  |  |
| --- | --- | --- | --- | --- | --- |
|  | dpi | W0 | W2 | W4 | W6 |
| **Untreated time course** | 3.19 ± 0.04 | 6.59 ± 0.10 |  |  |  |
| **Vehicle** |  |  | 6.40 ± 0.25 | 6.55 ± 0.22 | 6.42 ± 0.54 |
| **SnPP** |  |  | 5.52 ± 0.29 | 5.30 ± 0.30 | 5.30 ± 0.19 |
| **SnMP** |  |  | 5.73 ± 0.22 | 5.36 ± 0.31 | 5.98 ± 0.06 |
| **RHZ** |  |  |  | 4.17 ± 0.30 | 3.02 ± 0.30 |
| **RHZ + SnPP** |  |  |  | 4.11 ± 0.23 | 3.09 ± 0.16 |
| **RHZ + SnMP** |  |  |  | 4.01 ± 0.04 | 3.00 ± 0.12 |
| **SPaO** |  |  | 4.75 ± 0.17 | 3.18 ± 0.45 |  |
| **SPaO + SnPP** |  |  | 4.64 ± 0.13 | 2.67 ± 0.20 |  |
| **SPaO + SnMP5** |  |  | 4.68 ± 0.07 | 2.49 ± 0.25 |  |
| **Untreated relapse** | 2.52 ± 0.10 | 6.18 ± 0.11 |  |  |  |
| **SPaO** |  |  |  |  | 1.49 ± 0.27 |
| **SPaO + SnMP5** |  |  |  |  | 1.47 ± 0.30 |
| **SPaO + SnMP10** |  |  |  |  | 0.71 ± 0.23 |

### SnMP adjunctive therapy enhances the bactericidal efficacy of SPaO after 6 weeks of treatment without altering relapse rates

Next, we evaluated the adjunctive sterilizing activity of SnMP at different dosages (5 mg/kg and 10 mg/kg) in combination with SPaO. We selected treatment durations expected to yield > 50% relapse rates for the backbone regimen based on the prior literature [36]. The mice were treated for a total of 5 or 6 weeks, at which time the treatment was discontinued, and microbiological relapse was assessed 3 months later (Fig 4A). Adjunctive therapy with SnMP 5 mg/kg (SnMP5) did not significantly alter lung bacillary burdens relative to the SPaO regimen at the time points assessed. However, after 6 weeks of treatment, SPaO + SnMP 10 mg/kg (SnMP10) reduced mean lung bacterial burdens to 0.71 ± 0.23 log_10_ CFU, representing an additional 0.78 log_10_ CFU reduction (P = 0.005) when compared to SPaO alone (Fig 4B, Table 1). As in the bactericidal activity study, the lung/body weight ratios of mice treated with adjunctive SnMP5 or SnMP10 did not differ statistically from that of mice treated with the SPaO control regimen after 6 weeks of treatment (Fig S3A). Additionally, the lung/body weight ratios were maintained in each of the treatment groups at each relapse time point (Fig S3B). After 5 weeks of treatment, all mice experienced microbiological relapse with SPaO or SPaO + SnMP10 (Fig 4C). Similarly, the proportion of relapsing animals was equivalent in the groups receiving SPaO and SPaO + SnMP10 (66%) (Fig 4D).

**Figure 4:**
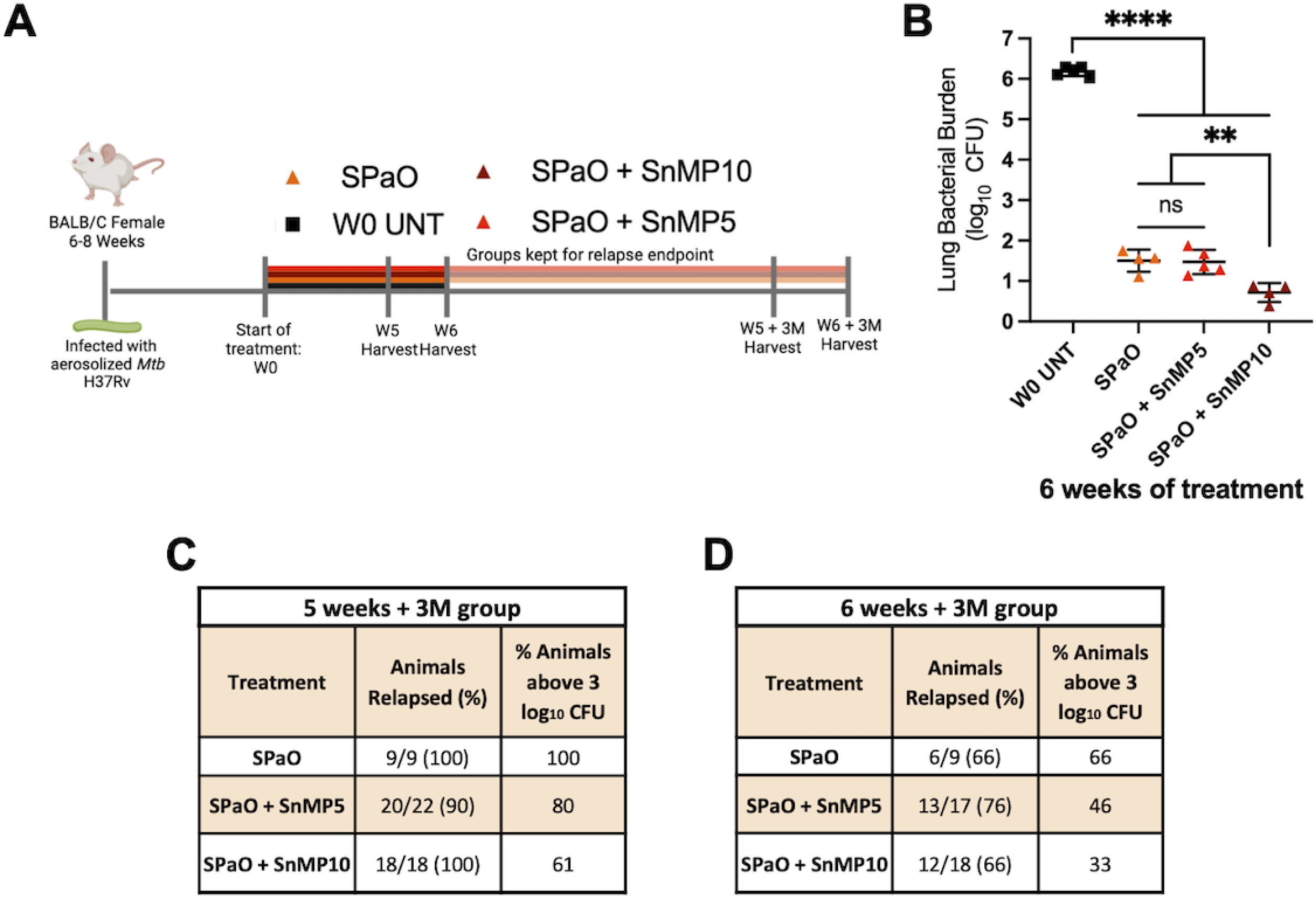
Adjunctive treatment with SnMP did not help reduce relapse rates after 6 weeks of treatment. A) Experimental design of mouse experiment. Each line represents a different treatment group and length of treatment. B) Lung bacillary burden after 6 weeks of treatment. Relapse study statistics containing the % relapsed animals, and animals above a 3 log_10_ CFU after completing 5 C) or 6 D) weeks of treatment and a three month wait period (+ 3M). W0= Week 0, W5= Week 5, W6= Week 6, SPaO: TBAJ-876(S)/Pretomanid(Pa)/TBI-223(O), Stannsoporfin: SnMP, SnMP5: SnMP 5 mg/kg, SnMP10: SnMP 10 mg/kg. Data in panel B represent four to five mice per treatment group. Statistical analysis was performed by an ordinary one-way ANOVA followed by Tukeys multiple comparison test. Data in panel C and D represent nine to twenty-two mice per treatment group. * = P < 0.01, ** = P < 0.001; **** = P < 0.0001, ns: not significant.

## Discussion

Inhibition of HO-1 has been proposed as a potential HDT strategy associated with reduced Mtb growth in vitro [37] and in vivo [20], [21]. Our study investigated the adjunctive bactericidal activity of a clinical-stage HO-1 inhibitor, SnMP, in a mouse model of chronic TB, as well as its adjunctive sterilizing activity when combined with an MDR-TB regimen in a murine model of microbiological relapse. We found that adjunctive therapy of SnMP enhanced the bactericidal activity of the SPaO regimen, but not that of the RHZ regimen, without exacerbating lung inflammation. Although adjunctive therapy with SnMP10 continued to enhance the bactericidal activity of SPaO after 6 weeks, it did not alter relapse rates compared to SPaO alone.

In contrast with prior work [20], we did not detect additive bactericidal activity of SnPP with RHZ in Mtb-infected mice. These discrepant findings may be explained by methodological differences in the two studies. In the current study, the administration of R was separated from that of HZ to limit drug interactions [38], [39], whereas the three antibiotics were administered together in the study by Costa *et al*. Furthermore, in the study by Costa *et al.*, C57BL/6 mice were used, while our study used BALB/c mice. Relative to BALB/c mice, C57BL/6 mice are able to more effectively limit the replication of Mtb through the development of T-cell-mediated immunity [40], [41]. Interestingly, this discrepancy was observed only when SnPP was given as adjunctive therapy, as we found that SnPP monotherapy reduced lung bacillary burden, as was reported previously [20]. These findings highlight the challenges associated with preclinical studies of HDT for TB, as the host-directed activity of these agents may be model-specific.

Our studies showed that SnMP administered alone regulated the expression of the gene encoding TNF-α in the lungs. Scharn *et al*. have shown previously that SnPP reduced the secretion of proinflammatory cytokines, including TNF-α, in Mtb-infected U937 cells [37], [24]. Notably, CD38 expression was upregulated in Mtb-infected mouse lungs following treatment with SnMP alone. Although CD38 is necessary for immune cell activation and is expressed by many immune cell types [42], it appears to be a specific marker for M1 macrophage phenotypes in murine models [35]. In humans and in mice, M1 macrophages play an essential role in propagating a host-protective response against Mtb (reviewed in Ahmad *et al.*, 2022) [43], [44]. Collectively, these results highlight an immunoregulatory role for the HO-1 inhibitor SnMP during Mtb infection. Additional studies are needed to further characterize the molecular mechanisms by which SnMP promotes Mtb clearance by the host.

Based on published data suggesting the SPaO regimen, at the doses of, S 12.5 mg/kg; Pa 100 mg/kg; and O 100 mg/kg, is able to eradicate lung infection in Mtb-infected mice by week 6 [36], the relapse time points of 5 and 6 weeks were selected to evaluate the adjunctive sterilizing activity of SnMP. In the current study, we found that, although SnMP increased the bactericidal activity of SPaO, adjunctive therapy did not reduce relapse rates compared to the background SPaO regimen. Future studies will focus on testing higher doses of SnMP and extending the duration of treatment to determine the potential of SnMP adjunctive therapy in shortening the duration of curative MDR treatment.

Consistent with prior preclinical studies [20], [21], [37], we have found that HO-1 inhibition is a promising HDT strategy for TB. We show for the first time that the novel HO-1 inhibitor, SnMP, enhances the bactericidal activity of the novel MDR-TB regimen SPaO in mice and modulates the expression of several pro-inflammatory cytokine and macrophage marker genes, which have been implicated in the control of Mtb replication in vivo, although adjunctive therapy with SnMP 10 mg/kg did not reduce relapse rates relative to SPaO. SnMP was well tolerated and did not alter gross lung pathology or histological inflammation in the lungs. Overall, this study advances our understanding of HO-1 inhibitors as potential adjunctive HDT agents for TB, but further research is needed to determine the potential utility of targeting this pathway against drug-susceptible and drug-resistant TB. In addition, to enhance the translational relevance of this HDT approach, future medicinal chemistry studies should focus on the development of formulations with favorable oral bioavailability and toxicity profiles.

## Materials and Methods

### Pharmacokinetic analyses

Single-dose PK studies were conducted by BioDuro Inc. (Beijing, China). SnMP and SnPP were formulated in sodium phosphate buffer (pH 7.4 – 7.8) and administered by the intraperitoneal route to 9–11-week-old female BALB/c mice. Three mice were used per each time point with 8 time points for each dose group. Blood was collected on 0.25, 0.5, 1, 2-, 4-, 8-, and 24-hours post-administration and levels of compounds in plasma were quantified by liquid chromatography-tandem mass spectrometry using AB Sciex API 4000 LC/MS/MS instrumentation. The following PK parameters were calculated from plasma drug concentrations: t1/2, tmax, Cmax, AUClast, AUCInf. All parameters were determined by noncompartmental analysis using WinNonLin software 8.0.

### Bacteria and growth conditions

Wild-type Mtb H37Rv was grown in Middlebrook 7H9 broth (Difco, Sparks, MD) supplemented with 10% oleic acid-albumin-dextrose-catalase (OADC) (Difco, Sparks, MD), 0.2% glycerol, and 0.05% Tween-80 at 37°C in a roller bottle.

### *M. tuberculosis* infection of mice

Wild-type Mtb H37Rv cultures were grown to an OD_600_ of 0.8-1, followed by 1:200 dilution in 7H9 broth (as stated above). 10 mL of the diluted bacterial culture was used as the inocolum for aerosol infection. Female BALB/c (6-8-week-old, Jackson Laboratory) were aerosol-infected with ∼100 bacilli of Mtb H37Rv using the Glas-Col Inhalation Exposure System (Terre Haute, IN). Animals were equally distributed in the five chambers of the Glas-Col machine to ensure equal infection amongst all animals. The inoculating culture was plated in serial tenfold dilutions on the day of infection on Middlebrook 7H11 agar containing 10% OADC. On the day post-infection, 5 mice were sacrificed, the lungs were homogenized using glass homogenizers and plated on 7H11 agar containing 10% OADC to enumerate the implanted CFU (Fig 1).

### Antibiotic and HDT treatment preparation

Protoporphyrin IX (SnPP) (Evotec, US) and stannsoporfin (SnMP) (Evotec, US) were prepared with sodium phosphate buffer (pH to 7.4-7.8 with HCl). Rifampin (R), isoniazid (H) and pyrazinamide (Z) (Sigma-Aldrich) were dissolved in deionized water. Pretomanid (Pa) (RTI) was dissolved in deionized water containing cyclodextrin micelle (CM-2) formulation,10% hydroxypropyl-β-cyclodextrin (HPCD) (Sigma) and 10% lecithin (ICN Pharmaceuticals Inc., Aurora, OH); TBAJ-876 (S) (Evotec, US) was prepared in 20% HPCD solution acidified with 1.5% 1N HCl, pH 3; and TBI-223 (O) (Evotec, US) was prepared in 0.5% Methylcellulose. The vehicle administered was either CM-2 formulation containing 10% HPCD (Sigma) and 10% lecithin (ICN Pharmaceuticals Inc., Aurora, OH) to mock Pa, 0.5% methylcellulose to mock O or 20% HPCD solution acidified with 1.5% 1N HCl to mock S. Vehicle of drugs were made in big batches and the drugs were dissolved into their corresponding vehicles weekly.

### Antibiotic and HDT doses and administration

In the bactericidal activity studies, the mice received one of the following regimens beginning 28 days after aerosol infection: 1) Vehicle (negative control); 2) SnPP 10 mg/kg; 3) SnMP 5 mg/kg; 4) R 10 mg/kg, H 25 mg/kg, Z 150 mg/kg (collectively, RHZ); 5) S 6.25 mg/kg, Pa 30 mg/kg and O 45 mg/kg (collectively, SPaO); 6) R 10 mg/kg, H 25 mg/kg, P 150 mg/kg, SnPP 10 mg/kg; 7) R 10 mg/kg, H 25 mg/kg, P 150 mg/kg, SnMP 5 mg/kg; 8) S 6.25 mg/kg, Pa 30 mg/kg, O 45 mg/kg, SnPP 10 mg/kg; 9) S 6.25 mg/kg, Pa 30 mg/kg, O 45 mg/kg, SnMP 5 mg/kg. The HO-1 inhibitors were given once daily by intraperitoneal injection (IP). PK analysis of the HO-1 inhibitors was conducted at 5 mg/kg and 10 mg/kg to test which dosages achieved similar plasma concentrations. There results also confirmed that the dosages achieved human-equivalent doses [45]. The other drugs were given orally by esophageal cannulation once daily, except for Pa and O, which were given twice daily separated by at least 4 hours. For the RHZ regimen, R was given first, and two hours later, H and Z were administered. The vehicle was administered via gavage to control mice concurrently with treatments containing active compounds to experimental mice.

In the relapse model studies, the mice received one of the following regimens beginning 28 days after aerosol infection: 1) Vehicle (negative control); 2) S 6.25 mg/kg, Pa 30 mg/kg, O 45 mg/kg; 3) S 6.25 mg/kg, Pa 30 mg/kg, O 45 mg/kg, SnMP 5 mg/kg; or 4) S 6.25 mg/kg, Pa 30 mg/kg, O 45 mg/kg, SnMP 10mg/kg. The discrepancies in numbers of animals between relapse groups is due to unexpected mortality from excessive gavage volume when the drugs were prepared separately at the start of therapy. Beginning in week 2, 2X concentrations of Pa and O were made and, at the time of administration, Pa and O were mixed at equal volumes to make a single gavage volume of 0.2 mL, which was tolerated well by the mice.

For both studies (except for the relapse time points, for which both lobes of the lung were homogenized and plated), the right lung and 3/5 of the left lung of each mouse were homogenized using glass homogenizers, and serial tenfold dilutions of lung homogenates in PBS were plated on Middlebrook 7H11 agar containing 0.4% activated charcoal to reduce the effects of drug carryover at the indicated time points; entire lung homogenates from relapse animals were plated on charcoal-containing 7H11 agar for consistency. Plates were incubated at 37°C and CFU were counted 4 weeks later. The CFU per lung data were calculated using lung weights, which were measured at the time of organ harvest. For the animals in the relapse groups, the entire lung was used for CFU determination. Microbiological relapse was defined as the presence of 1 or more colonies in plated lung homogenates. All procedures were performed according to protocols approved by the Johns Hopkins University Institutional Animal Care and Use Committee.

### Measurement of lung/body weight ratio

Lung and body weight measurements in milligrams (mg) and grams (g), respectively, were taken at the time of harvest using weight scales. The lung/body weight ratio was calculated by dividing lung(g)/body(g) weight and then multiplying by 100.

### Quantitative analysis of lung histopathology

At the time of harvest, the left lung medial apex to base was sectioned and fixed with 4% paraformaldehyde (PFA). After a 24-hour incubation in 4% PFA, the lung section was transferred to PBS and sent in for processing. The paraffin blocks of lung samples were processed to obtain three sections per lung spaced 12 μm apart for hematoxylin and eosin staining. Each lung section was sectioned to 4 μm. Each slide was scanned by a brightfield microscope at the JHU Oncology Tissue Core Center to obtain digital 40X images. Quantitative analysis was performed by JRC and AY blinded to treatment allocation and mouse strain. For each treatment group, a total of 15 scanned slides (3 slides per mice) were reviewed. Regions of interest corresponding to the absolute areas of total lung surface area and inflammation were manually drawn and their area quantified using the open-source software QuPath (https://qupath.github.io/). Inflammation was defined as: 1. “Lymphocytic cuffs”, areas of high cell density due to the predominance of lymphocytes, which are smaller cells with scant cytoplasm; 2. Regions dense with macrophages, which have relatively large amounts of cytoplasm. The lung inflammation to whole surface area was quantified and multiplied by 100 to obtain the percentage.

### Cytokine or cell marker gene expression

A section of the left lower lobe (∼ 1/5 of the left lung) was flash-frozen in liquid nitrogen at the time of harvest and later lysed in TRIzol using a hard tissue homogenizing kit (CK28-R, Percellys) with the manufacturer-recommended bead-beating protocol (Percellys Evolution; 8,500 rpm, three 15-s cycles with 30-s breaks on ice after each cycle). Total RNA was extracted using the miRNeasy minikit (217004, Qiagen). RT-qPCR sequences for each primer pair were ordered from IDT and are listed in the supplementary material Table 2. Two-step RT-qPCR was performed on a StepOnePlus real-time PCR system (ThermoFisher Scientific) using the high-capacity RNA-to-cDNA kit (4387406, ThermoFisher Scientific), followed by Power SYBR green PCR master mix (436877, Applied Biosystems). All samples were tested with three technical replicates. Target gene expression (cycle threshold (Ct) values) were normalized relative to the housekeeping gene *β-actin*. The Ct value for each gene in infected test samples was then normalized to that of uninfected samples and mRNA expression fold change was determined by the ΔΔCt method.

**Table 2:**
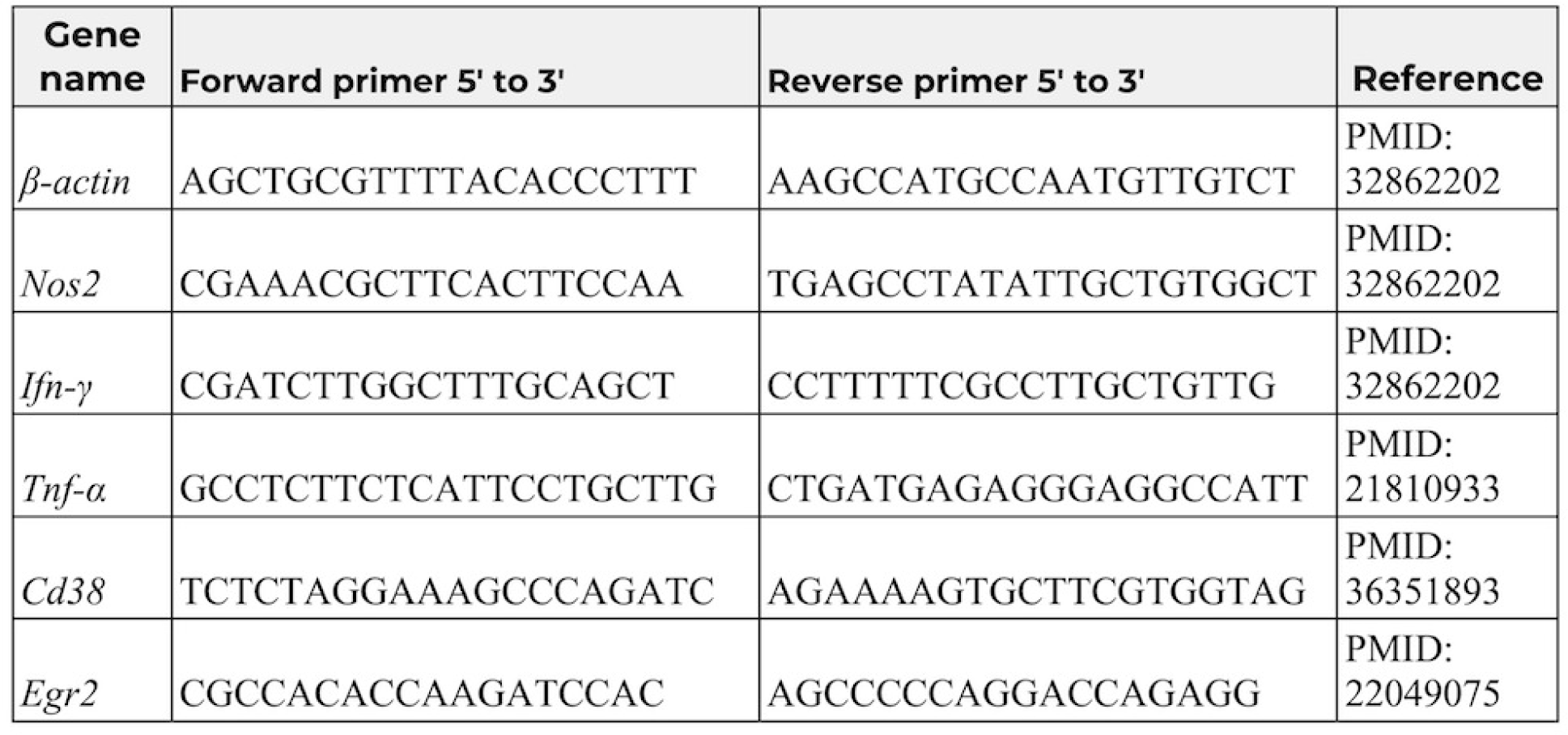
RT-qPCR sequences for each primer pair.

### Statistics

The differences between groups were assessed using an ordinary one-way ANOVA followed by different multiple comparison tests or Welch’s ANOVA test (depending on the spread of the data) as stated on the figure panel description. The Prism software (GraphPad, San Diego, CA, USA) version 9 was utilized for this analysis. Results were considered statistically significant when the p-value was less than 0.05. The Pearson’s correlation coefficient was calculated using the Prism software.

## Supplemental material

**Figure S1:** Plasma pharmacokinetic parameters for SnPP and SnMP in mice.

**Figure S2:** Lung/body weight ratio correlates with lung bacterial burden after 2, 4 and 6 weeks of treatment.

**Figure S3:** Mouse lung/body weight ratio after 6 weeks of treatment and at the relapse time point.

**Table 1:** Lung CFU count for time course and relapse study.

**Table 2:** RT-qPCR sequences for each primer pair.

## Author Contributions

PN, SKa, NS, and PCK conceived and designed the time course study. NS provided data and the description for the pharmokinetic study. PN, SKa, SA administered the therapies; PN, SKa, SKr, performed animal haversting, tissue sectioning and weight collection, imaged lungs, plated the homogenates of the infected lungs, collected CFU and analysed CFU data for the time course study. JRC performed data analysis for the lung/body weight ratio, and the lung/body weight ratio vs CFU correlation. JRC & AY performed the quantitative histology analysis. JRC extracted RNA from lung tissue. JRC and DQ performed RTqPCR experiements, and performed data analysis. JRC, PN, Ska, NS, and PCK conceived and designed the relapse study. JRC, and PN administered the therapies for the relapse study. JRC, PN, and Ska performed animal haversting, tissue sectioning and weight collection; PN, JRC, and AG plated the homogenates of the infected lungs; JRC, PN performed CFU data collection and analysis for the relapse study. JRC, and PCK drafted the manuscript. JRC, PN, Ska, SKr, DQ, AG, SA, AY, DLC, AS, NF, NS, and PCK interpreted the data and edited the manuscript. All authors edited and approved the final manuscript.

## Acknowledgments

These studies were supported by NIAID/NIH grant K24AI143447 and a grant from the TB Alliance to PCK. AS is supported by the intramural research program of the NIAID. We thank the Oncology Tissue Services (SKCCC) at JHU supported by the P30 CA006973 grant for their services and for imaging the H&E-stained slides. We gratefully acknowledge the assistance of Michael Urbanowski, MD, PhD, for his training and guidance in the histopathological quantification of the lung tissues.

## Ethics statement

The animal study was reviewed and approved by Johns Hopkins University Institutional Animal Care and Use Committee. The animal welfare assurance number is D16-00173 (A3272-01). JHU is registered with the USDA to conduct animal research and has maintained active AAALAC accreditation since 10/4/1974.

## References

[1] World Health Organization, “Global Tuberculosis Report 2022,” 2022. [Online]. Available: http://apps.who.int/bookorders.

[2] K. J. Seung, S. Keshavjee, and M. L. Rich, “Multidrug-resistant tuberculosis and extensively drug-resistant tuberculosis,” Cold Spring Harb Perspect Med, vol. 5, no. 9, Sep. 2015, doi: 10.1101/cshperspect.a017863.

[3] E. Pontali et al., “Regimens to treat multidrug-resistant tuberculosis: Past, present and future perspectives,” European Respiratory Review, vol. 28, no. 152. European Respiratory Society, 2019. doi: 10.1183/16000617.0035-2019.

[4] C. Lange, D. Chesov, J. Heyckendorf, C. C. Leung, Z. Udwadia, and K. Dheda, “Drug-resistant tuberculosis: An update on disease burden, diagnosis and treatment,” Respirology, vol. 23, no. 7, pp. 656–673, Jul. 2018, doi: 10.1111/resp.13304.

[5] D. J. Frank et al., “Remembering the host in tuberculosis drug development,” in *Journal of Infectious Diseases*, Oxford University Press, May 2019, pp. 1518–1524. doi: 10.1093/infdis/jiy712.

[6] G. Kilinç, A. Saris, T. H. M. Ottenhoff, and M. C. Haks, “Host-directed therapy to combat mycobacterial infections*,” Immunological Reviews, vol. 301, no. 1. John Wiley and Sons Inc, pp. 62–83, May 01, 2021. doi: 10.1111/imr.12951.

[7] R. Tenhunen, H. S. Marver, and R. Schmid, “The enzymatic conversion of heme to bilirubin by microsomal heme oxygenase.,” Proc Natl Acad Sci U S A, vol. 61, no. 2, pp. 748–55, Oct. 1968, doi: 10.1073/pnas.61.2.748.

[8] S. W. Ryter and A. M. K. Choi, “Heme oxygenase-1/carbon monoxide: From metabolism to molecular therapy,” American Journal of Respiratory Cell and Molecular Biology, vol. 41, no. 3. pp. 251–260, Sep. 01, 2009. doi: 10.1165/rcmb.2009-0170TR.

[9] M. D. Maines, “THE HEME OXYGENASE SYSTEM: A Regulator of Second Messenger Gases,” 1997. [Online]. Available: www.annualreviews.org

[10] S. W. Ryter and R. M. Tyrrell, “The heme synthesis and degradation pathways: role in oxidant sensitivity. Heme oxygenase has both pro- and antioxidant properties.,” Free Radic Biol Med, vol. 28, no. 2, pp. 289–309, Jan. 2000, doi: 10.1016/s0891-5849(99)00223-3.

[11] N. Rockwood et al., “Mycobacterium tuberculosis induction of heme oxygenase-1 expression is dependent on oxidative stress and reflects treatment outcomes,” Front Immunol, vol. 8, no. MAY, May 2017, doi: 10.3389/fimmu.2017.00542.

[12] A. Kumar, A. Farhana, L. Guidry, V. Saini, M. Hondalus, and A. J. C. Steyn, “Redox homeostasis in mycobacteria: the key to tuberculosis control?,” Expert reviews in molecular medicine, vol. 13. 2011. doi: 10.1017/s1462399411002079.

[13] M. H. Kapturczak et al., “Animal Model Heme Oxygenase-1 Modulates Early Inflammatory Responses Evidence from the Heme Oxygenase-1-Deficient Mouse,” 2004.

[14] B. Wegiel et al., “Carbon monoxide expedites metabolic exhaustion to inhibit tumor growth,” Cancer Res, vol. 73, no. 23, pp. 7009–7021, Dec. 2013, doi: 10.1158/0008-5472.CAN-13-1075.

[15] L. E. Otterbein et al., “Carbon monoxide has anti-inflammatory effects involving the mitogen-activated protein kinase pathway,” Nat Med, vol. 6, no. 4, pp. 422–428, Apr. 2000, doi: 10.1038/74680.

[16] M. P. Soares and F. H. Bach, “Heme oxygenase-1: from biology to therapeutic potential,” Trends Mol Med, vol. 15, no. 2, pp. 50–58, Feb. 2009, doi: 10.1016/j.molmed.2008.12.004.

[17] Y. Naito, T. Takagi, and Y. Higashimura, “Heme oxygenase-1 and anti-inflammatory M2 macrophages,” Archives of Biochemistry and Biophysics, vol. 564. Academic Press Inc., pp. 83–88, Dec. 15, 2014. doi: 10.1016/j.abb.2014.09.005.

[18] S. C. Funes, M. Rios, J. Escobar-Vera, and A. M. Kalergis, “Implications of macrophage polarization in autoimmunity,” Immunology, vol. 154, no. 2. Blackwell Publishing Ltd, pp. 186–195, Jun. 01, 2018. doi: 10.1111/imm.12910.

[19] S. O’Leary, M. P. O’Sullivan, and J. Keane, “IL-10 blocks phagosome maturation in Mycobacterium tuberculosis-infected human macrophages,” Am J Respir Cell Mol Biol, vol. 45, no. 1, pp. 172–180, Jul. 2011, doi: 10.1165/rcmb.2010-0319OC.

[20] D. L. Costa et al., “Pharmacological inhibition of host heme oxygenase-1 suppresses mycobacterium tuberculosis infection in vivo by a mechanism dependent on T lymphocytes,” mBio, vol. 7, no. 5, Sep. 2016, doi: 10.1128/mBio.01675-16.

[21] D. L. Costa et al., “Heme oxygenase-1 inhibition promotes IFNγ- and NOS2-mediated control of Mycobacterium tuberculosis infection,” Mucosal Immunol, vol. 14, no. 1, pp. 253–266, Jan. 2021, doi: 10.1038/s41385-020-00342-x.

[22] American Academy of Pediatrics Subcommittee on Hyperbilirubinemia, “Management of hyperbilirubinemia in the newborn infant 35 or more weeks of gestation.,” Pediatrics, vol. 114, no. 1, pp. 297–316, Jul. 2004, doi: 10.1542/peds.114.1.297.

[23] T. D. Burt, L. Seu, J. E. Mold, A. Kappas, and J. M. McCune, “Naive Human T Cells Are Activated and Proliferate in Response to the Heme Oxygenase-1 Inhibitor Tin Mesoporphyrin,” The Journal of Immunology, vol. 185, no. 9, pp. 5279–5288, Nov. 2010, doi: 10.4049/jimmunol.0903127.

[24] B. W. Allwood et al., “Post-tuberculosis lung health: Perspectives from the First International Symposium,” in *International Journal of Tuberculosis and Lung Disease*, International Union Against Tuberculosis and Lung Disease (The Union), Aug. 2020, pp. 820–828. doi: 10.5588/ijtld.20.0067.

[25] J. G. Pasipanodya et al., “Pulmonary impairment after tuberculosis and its contribution to TB burden,” BMC Public Health, vol. 10, 2010, doi: 10.1186/1471-2458-10-259.

[26] F. J. Leong, V. Dartois, and T. Dick, Eds., A Color Atlas of Comparative Pathology of Pulmonary Tuberculosis, 1st Edition. CRC Press, 2010. doi: 10.1201/EBK1439835272.

[27] K. D. Poss and S. Tonegawa, “Heme oxygenase 1 is required for mammalian iron reutilization,” 1997. [Online]. Available: www.pnas.org.

[28] J. P. Juncos et al., “Anomalous renal effects of tin protoporphyrin in a murine model of sickle cell disease,” American Journal of Pathology, vol. 169, no. 1, pp. 21–31, 2006, doi: 10.2353/ajpath.2006.051195.

[29] C. E. Bunse et al., “Modulation of heme oxygenase-1 by metalloporphyrins increases anti-viral T cell responses,” Clin Exp Immunol, vol. 179, no. 2, pp. 265–276, Feb. 2015, doi: 10.1111/cei.12451.

[30] J. M. Cicchese et al., “Dynamic balance of pro- and anti-inflammatory signals controls disease and limits pathology,” Immunol Rev, vol. 285, no. 1, pp. 147–167, Sep. 2018, doi: 10.1111/imr.12671.

[31] M. K. K. Niazi, G. Beamer, and M. N. Gurcan, “Detecting and characterizing cellular responses to Mycobacterium tuberculosis from histology slides.,” Cytometry A, vol. 85, no. 2, pp. 151–61, Feb. 2014, doi: 10.1002/cyto.a.22424.

[32] N. V Serbina and J. L. Flynn, “Early Emergence of CD8 T Cells Primed for Production of Type 1 Cytokines in the Lungs of Mycobacterium tuberculosis-Infected Mice,” 1999.

[33] I. E. A. Flesch and S. H. E. Kaufmann, “Activation of Tuberculostatic Macrophage Functions by Gamma Interferon, Interleukin-4, and Tumor Necrosis Factor,” 1990.

[34] M. Safford et al., “Egr-2 and Egr-3 are negative regulators of T cell activation,” Nat Immunol, vol. 6, no. 5, pp. 472–480, 2005, doi: 10.1038/ni1193.

[35] K. A. Jablonski et al., “Novel markers to delineate murine M1 and M2 macrophages,” PLoS One, vol. 10, no. 12, Dec. 2015, doi: 10.1371/journal.pone.0145342.

[36] S. Y. Li et al., “Next-Generation Diarylquinolines Improve Sterilizing Activity of Regimens with Pretomanid and the Novel Oxazolidinone TBI-223 in a Mouse Tuberculosis Model,” Antimicrob Agents Chemother, vol. 67, no. 4, Apr. 2023, doi: 10.1128/aac.00035-23.

[37] C. R. Scharn et al., “ Heme Oxygenase-1 Regulates Inflammation and Mycobacterial Survival in Human Macrophages during Mycobacterium tuberculosis Infection,” The Journal of Immunology, vol. 196, no. 11, pp. 4641–4649, Jun. 2016, doi: 10.4049/jimmunol.1500434.

[38] J. Grosset, C. Truffot-Pernot, C. Lacroix, and B. Ji, “Antagonism between isoniazid and the combination pyrazinamide-rifampin against tuberculosis infection in mice.,” Antimicrob Agents Chemother, vol. 36, no. 3, pp. 548–51, Mar. 1992, doi: 10.1128/AAC.36.3.548.

[39] Z. Ahmad et al., “Comparison of the ‘Denver regimen’ against acute tuberculosis in the mouse and guinea pig.,” J Antimicrob Chemother, vol. 65, no. 4, pp. 729–34, Apr. 2010, doi: 10.1093/jac/dkq007.

[40] A. M. Cooper, “Cell-mediated immune responses in tuberculosis,” Annual Review of Immunology, vol. 27. pp. 393–422, 2009. doi: 10.1146/annurev.immunol.021908.132703.

[41] J. L. Flynn and J. Chan, “Immunology of tuberculosis.,” Annu Rev Immunol, vol. 19, pp.93–129, 2001, doi: 10.1146/annurev.immunol.19.1.93.

[42] W. Li et al., “CD38: A Significant Regulator of Macrophage Function,” Frontiers in Oncology, vol. 12. Frontiers Media S.A., Feb. 17, 2022. doi: 10.3389/fonc.2022.775649.

[43] F. Ahmad et al., “Macrophage: A Cell With Many Faces and Functions in Tuberculosis,” Frontiers in Immunology, vol. 13. Frontiers Media S.A., May 06, 2022. doi: 10.3389/fimmu.2022.747799.

[44] A. Khan et al., “Human M1 macrophages express unique innate immune response genes after mycobacterial infection to defend against tuberculosis,” Commun Biol, vol. 5, no. 1, Dec. 2022, doi: 10.1038/s42003-022-03387-9.

[45] Food and Drug Administration and GASTROINTESTINAL DRUGS ADVISORY COMMITTEE and the PEDIATRIC ADVISORY COMMITTEE, “Stannsoporfin - Proposed for the treatment of neonates greater than or equal to 35 weeks of gestational age with indicators of hemolysis who are at risk of developing severe hyperbilirubinemia,” May 2018. Accessed: Aug. 01, 2023. [Online]. Available: https://fda.report/media/112936/FDA-Briefing-Information-for-the-May-3--2018-Joint-Meeting-of-the-Gastrointestinal-Drugs-Advisory-Committee-and-the-Pediatric-Advisory-Committee.pdf

