## Supplemental figs S1-3 for "The heme oxygenase-1 metalloporphyrin inhibitor stannsoporfin enhances the bactericidal activity of a novel regimen for multidrug-resistant tuberculosis in a murine model": Fig S1.pdf

A

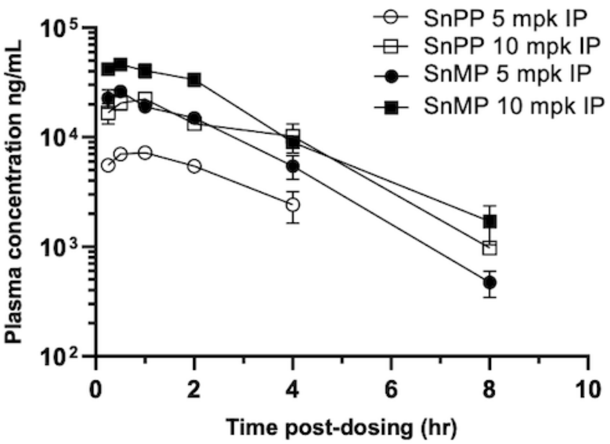


B

| PK parameters | SnPP 5 mpk | SnPP 10 mpk | SnMP 5 mpk | SnMP 10 mpk |
|---|---|---|---|---|
| $C_{max}$ (ng/mL) | 7523 | 22533 | 37179 | 64117 |
| $AUC_{last}$ (hr*ng/mL) | 20058 | 81085 | 88030 | 185683 |
| $T_{1/2}$ (hr) | 1.97 | 1.52 | 1.3 | 1.4 |

**Figure S1: Plasma pharmacokinetic parameters for SnPP and SnMP in mice.**

9–11-week-old female BALB/c mice were administered with SnPP or SnMP by intraperitoneal route. Blood was collected on 0.25, 0.5, 1, 2-, 4-, 8-, and 24-hours post-administration and levels of compounds in plasma were quantified by liquid chromatography-tandem mass spectrometry using AB Sciex API 4000 LC/MS/MS instrumentation A) Plasma concentrations (ng/mL) against time post-dosing up to 8 hours. B) Summary of pharmacokinetic parameters for SnPP and SnMP. PK parameters were calculated from plasma drug concentrations: t1/2, tmax, Cmax, AUClast, AUCInf. All parameters were determined using were determined by noncompartmental analysis using WinNonLin software 8.0.
