## Supplemental figs S1-3 for "The heme oxygenase-1 metalloporphyrin inhibitor stannsoporfin enhances the bactericidal activity of a novel regimen for multidrug-resistant tuberculosis in a murine model": Fig S2.pdf

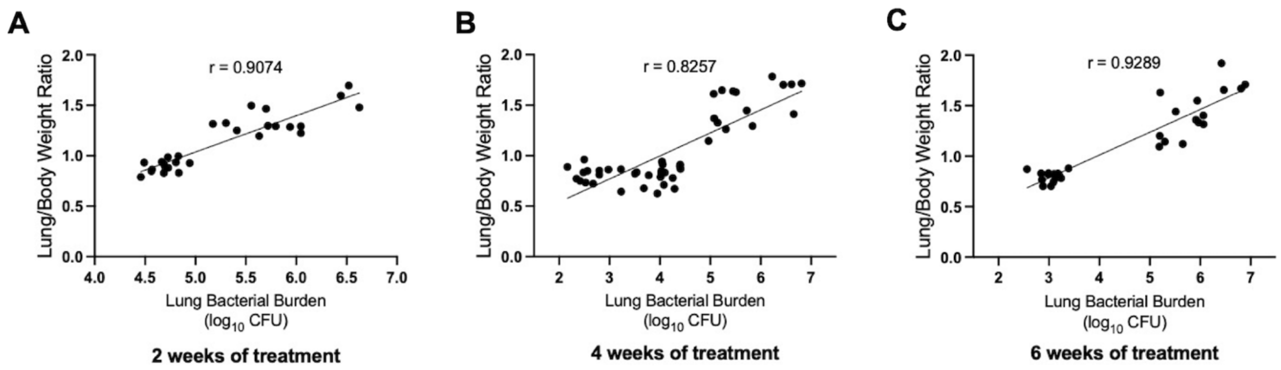


**Figure S2: Lung/body weight ratio correlates with lung bacterial burden after 2, 4 and 6 weeks of treatment.**

Scatterplot of matched animal lung/body weight ratio and lung bacterial burden after 2 A), 4 B), and 6 C) weeks of treatment. Each dot represents one animal. r= Pearson's correlation coefficient.
