## Supplemental figs S1-3 for "The heme oxygenase-1 metalloporphyrin inhibitor stannsoporfin enhances the bactericidal activity of a novel regimen for multidrug-resistant tuberculosis in a murine model": Fig S3.pdf

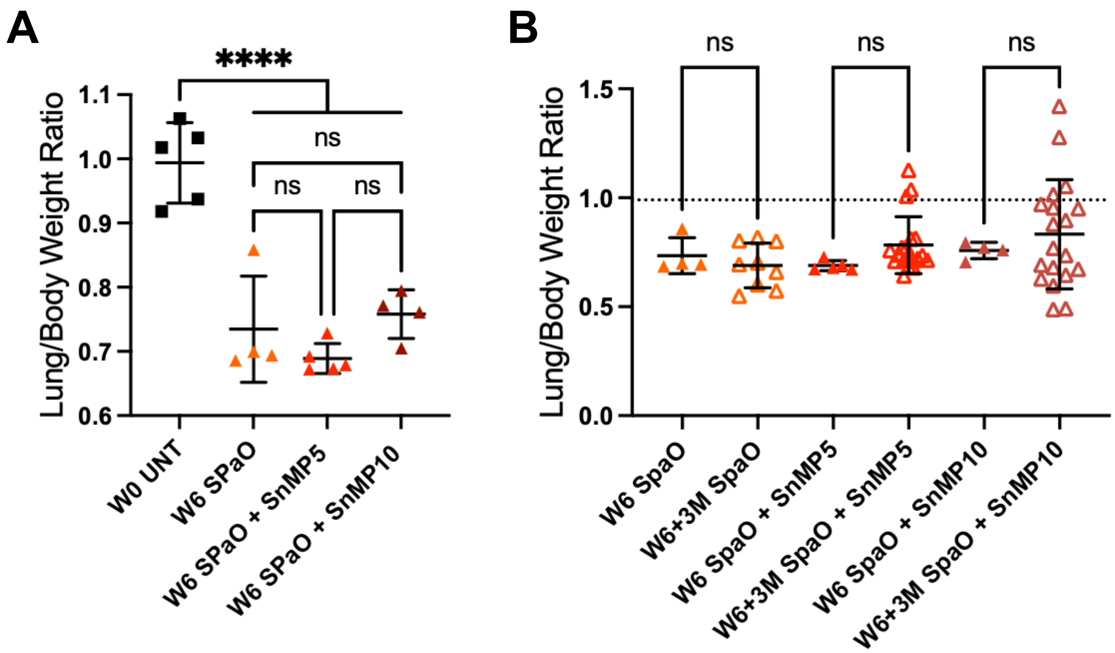


**Figure S3: Mice lung/body weight ratio after 6 weeks of treatment and at the relapse timepoint.**

A, B) Lung and body weights were taken at time of harvest B) The dotted line indicates the average whole lung weight for the W0 UNT harvest group. Data show means ± SD of the lung weight to body weight ratio. W0= Week 0, W6= Week 6, W6+3M= Week 6 plus 3 months, UNT= untreated, SPaO: TBAJ-876(S)/Pretomanid(Pa)/TBI-223(O), Stannsoporfin: SnMP, SnMP5: SnMP 5 mg/kg, SnMP10: SnMP 10 mg/kg. Panel A had four to five mice per treatment group, panel B had four to twenty-two mice per treatment group. Statistical analysis was performed by a one-way ANOVA followed by Tukeys multiple comparison test for panels A and B. **** = P < 0.0001, ns = not significant.
